# Anti-fungal recombinant psoriasin effectively inhibits Candida albicans growth on denture base

**DOI:** 10.1101/2024.03.12.584579

**Authors:** Lucia Adriana Lifshits, Edward Brohnshtein, May Attias, Yoav Breuer, Adi Cohen, Matan Gabay, Marina Sova, Evgeny Weinberg, Eran Zenziper, Daniel Z. Bar, Nir Sterer, Maayan Gal

## Abstract

Oral candidiasis leading to denture stomatitis is a fungal infection resulting from unregulated growth and adhesion mainly of *Candida albicans* onto acrylic denture base. Once the biofilm is formed, it is immune resistant and mainstay treatments involve toxic chemical antifungal agents or mechanical cleaning techniques, both offer limited efficacy. Consequently, there is an urgent need for more effective and safer therapeutic approaches. While biological modalities are expanding in general medicine, the exploration of protein-based therapeutics in dental medicine remains limited. This research evaluates the inhibitory effect of recombinantly expressed psoriasin on the growth of *Candida albicans* on polymethyl methacrylate denture bases. Psoriasin, also known as S100-A7, has shown promise in treating microbial skin infections, and its natural presence in saliva makes it a promising candidate for treating oral microbial infections. Our findings indicate that psoriasin exhibits a strong, dose-dependent inhibition of *Candida albicans* growth. Further, we incubated a polymethyl methacrylate denture base within the psoriasin solution. Notably, immersing the denture base in the solution completely eradicated fungal growth. Our research utilizes natural antifungal proteins within biomedical devices like denture bases, suggesting psoriasin as a safe alternative to chemical antifungals in dental medicine.

## 1. Introduction

Oral candidiasis, predominantly caused by *Candida species*, is a common human fungal infection(1-3). Denture stomatitis (chronic atrophic candidiasis) resulting from the adhesion and growth of *Candida albicans* (*C. albicans*) onto the denture’s acrylic base(4-6), impacts approximately 65% of denture wearers(1,7,8), and poses a particular risk to immunocompromised patients(9). This is of particular importance in an elderly population, where dentures are commonly used, and poor oral hygiene, together with a declined immune system, leads to a high rate of acute candidiasis. Denture stomatitis may evolve into pre-cancerous lesions like candida leukoplakia(10). Moreover, biofilm formation affects the mucosa by inducing a painful inflammation in the tissues adjacent to the contaminated dentures(11,12). It is thus important to address this widespread oral health issue.

Dentures are primarily fabricated from polymethyl methacrylate (PMMA), an acrylic base material. The inherent porosity of PMMA, resulting from air entrapment during its manufacturing, makes it ideal for microbial adhesion and biofilm formation(13,14). Specifically, *Candida* species have a high affinity for adhering to PMMA-based dentures. This adhesion often leads to the transformation of *Candida* into a more virulent form, promoting biofilm formation, which is a significant concern in dental prosthetics(2,3).

Biofilms on dentures show resistance to host immunity and are challenging to eliminate without mechanical cleaning or harsh chemical agents. Indeed, current treatments include physical cleaning and antifungal agents like Amphotericin B, Azoles, and Nystatin(15-18). However, prolonged use of these chemicals can lead to resistance and adverse toxic side effects(19,20). Additional preventive strategies that aim to overcome the limitations of current treatments include the incorporation of antimicrobial nanoparticles in the denture base(21,22), the use of alternative denture fabrication techniques(23), the application of natural plant extracts(24), and medium-chain fatty acids(25).

The objective of this study is to explore an alternative strategy for prevention and treatment of denture stomatitis by applying natural anti-fungal proteins that are found in the human saliva(26). Among these, psoriasin (S100-A7) is a member of the S100 calcium-binding protein family known for its role in immune responses and wound healing(27). Psoriasin has effective antibacterial and antifungal properties(28), and studies have shown it can disrupt fungal cell membranes, leading to fungal death(29). Moreover, previous studies have shown psoriasin has an effective antimicrobial activity(26,30). Our research shows that recombinantly expressed psoriasin effectively inhibits *C. albicans* growth on PMMA-based dentures and is safe for human cell lines, highlighting its potential as a novel treatment modality for denture stomatitis.

## 2. Materials and Methods

### 2.1 Expression and purification of recombinant psoriasin

The psoriasin gene, optimized for *E. coli* expression with an N-terminal HISx6 tag, was synthesized and cloned into the pET-28a vector (Genscript Ltd). The plasmid was transformed into *E. coli* BL21 (DE3) pLysS. Bacterial cultures were grown at 37°C and 200 rpm until an OD600 of ∼0.8 was reached. Protein expression was induced with 1 mM IPTG for 16 hours at 25°C. Following sonication, the lysate was centrifuged at 16,000xg for 30 minutes. The supernatant was loaded onto nickel column chromatography, and following extensive washing, the protein was subsequently eluted using 300 mM imidazole. The protein was then buffer-exchanged into PBS and frozen with 50% glycerol. For the analysis of the protein on SDS-PAGE gel, samples from each purification step were collected and resuspended in 4X Laemmli sample buffer (BioRad, Cat. #1610747) and boiled at 95°C for 10 min. Samples were subjected to 12% SDS-PAGE gel and stained with Coomassie brilliant blue (Sigma-Aldrich, Cat. #115444).

### 2.2 *C. albicans* growth curves and absorbance

To monitor *C. albicans* growth, 10 µl from its glycerol stock was diluted in 10 ml of RPMI-MOPS and recovered at 30°C for 12 hours. Thereafter, the optical density at 600 nm (OD_600_) was measured and adjusted to an initial optical density of 0.001 by diluting the stock into a fresh RPMI-MOPS. Subsequently, 100 µL aliquots were transferred to a transparent 96-well plate and mixed with the indicated psoriasin concentrations. The plate was incubated in a Biotek Synergy H1 microplate reader at 30°C under continuous orbital shaking. Absorbance readings were taken at 600 nm at 10-minute intervals from three replicate wells.

### 2.3 PMMA discs

PMMA denture base acrylic discs, each with a diameter of 12mm and a thickness of 2mm, were fabricated via hot curing according to the manufacturer’s instructions (Novodon, Novodent ETS).

### 2.4 Soaking of PMMA discs with *C. albicans*

PMMA discs were placed in 50 mL tubes filled with either 10 mL of PBS, PBS supplemented with 50 µM psoriasin or 28 µM voriconazole. After a 24-hour incubation, all discs were rinsed with PBS and transferred to a 24-well plate containing 1 mL RPMI-MOPS inoculated with *C. albicans*. The plate was subjected to orbital shaking at 30°C for 24 hours. Results show the average of three replicate discs.

### 2.5 Microscopy images

Images were captured at 20x magnification in brightfield mode (Bio-Rad ZOE cell imager).

### 2.6 PDB structure of psoriasin

The structure of human S100A7 was obtained from the Protein Data Bank (PDB ID: 4AQJ) and visualized using ChimeraX(31).

### 2.7 Cellular viability

Experiments involving primary human gingival fibroblasts (hGFs) received approval from the Tel Aviv University Institutional Review Board (IRB No. 0001006-1). Informed consent was acquired from all participating patients. Both HeLa and hGFs were cultured as described earlier(32,33) in Dulbecco’s modified Eagle’s medium (DMEM), containing 0.11 g/liter sodium pyruvate, 2 mM l-glutamine, 4.5 g/liter glucose, 10% fetal bovine serum (FBS), 100 U/ml penicillin, and 100 U/ml streptomycin in a humidified incubator at 5% (v/v) CO_2_. Cells were seeded in a 96-well plate (5,000 cells per well) and incubated with variable concentrations of psoriasin. Cellular viability was evaluated by replacement of growth medium with DMEM containing 10% (v/v) of Alamar blue, and cells were further incubated for 3 hours at 37°C in humidified 95% air, 5% CO_2_. Alamar blue measurements were taken using a BioTek H1 synergy microplate reader. The absorbance of each sample was measured at an excitation/emission wavelength of 530/590 nm.

### 2.8 Scanning Electron Microscopy

Discs were fixed in 2.5% glutaraldehyde in PBS. After washing with PBS, samples were dehydrated by successive ethanol treatment. After critical point drying (Balzer’s critical point dryer), the discs were mounted on aluminum stubs and sputter-coated (SC7620, Quorum) with gold. Images were captured on the scanning electron microscope (JCM-6000, JEOL).

### 2.9 Image analysis

Images analysis and quantification were done with Fiji/ImageJ (version 2.14.0/1.54f), https://imagej.net/software/fiji/downloads.

## 3. Results

### 3.1 Psoriasin recombinant expression and purification

Relying upon recombinant proteins as therapeutic agents necessitates a requisite for their high-yield and robust expression. To this end, we established a recombinant expression and purification protocol of psoriasin in *E. coli*. We synthesized a codon-optimized gene for optimal expression in *E. coli*, containing an N-terminus Hisx6 suitable for large-scale expression and subsequent chromatography-based affinity-tag purification (Supplementary Figure 1). Despite the protein’s inherent antimicrobial activity including against *E. coli* (34), the tight regulation of its expression in the BL21 pLysS strain enables bacterial growth to the desired cells density and ensures adequate induction of protein expression. In non-induced bacteria, psoriasin’s basal expression was undetectable (Fig. 1, lane 2), while following IPTG induction it demonstrated high expression levels (Fig. 1, lanes 3-4). Nickel column purification effectively isolated the desired protein (Fig. 1, lanes 4-8).

**Figure 1.**
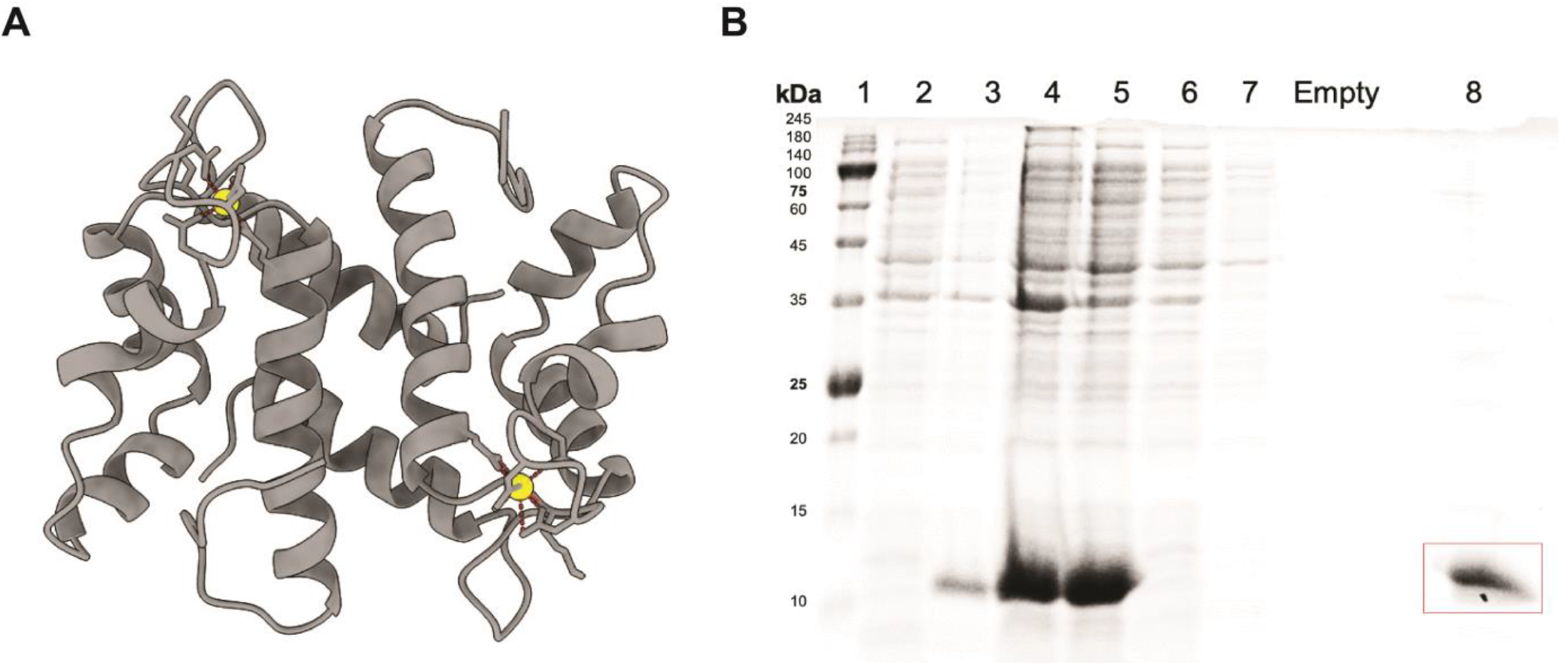
Psoriasin Expression and Purification. **(A)** Structure of human S100-A7 psoriasin dimer based on Protein Data Bank entry 4AQJ; calcium ions are depicted in yellow. **(B)** SDS-PAGE gel illustrating psoriasin purification. The *E. coli* BL21 lysate containing the protein was loaded onto nickel column; non-specific binders were washed prior to protein elution. Lanes: 1-Protein marker; 2-Uninduced cells; 3-4-Cells 3 and 16 hours post-IPTG induction; 5-Supernatant from cell lysate; 6-7-Column wash; 8-Protein eluted with 300 mM imidazole. The calculated molecular weight of the 6xHis-tagged psoriasin is 13.6 kDa.

### 3.2 Psoriasin inhibits *C. albicans* growth in a dose-dependent manner

Following the successful expression and purification of psoriasin (Fig. 1), we assessed its ability to inhibit *C. albicans* culture growing in a 96-well plate. Culturing *C. albicans* in RPMI-MOPS at 30°C with varying psoriasin concentrations revealed a dose-dependent inhibitory effect (Fig. 2A). Untreated cells (top green line) exhibited normal growth, however, psoriasin concentrations above 300 nM completely suppressed fungal growth. Figure 2B shows a bar plot analyzing the maximum absorbance at the end of the growth period for each treatment concentration. Fitting the data to a variable slope sigmoidal curve, resulting in an IC50 of 115 nM, confirming psoriasin’s efficacy under these experimental conditions.

**Figure 2.**
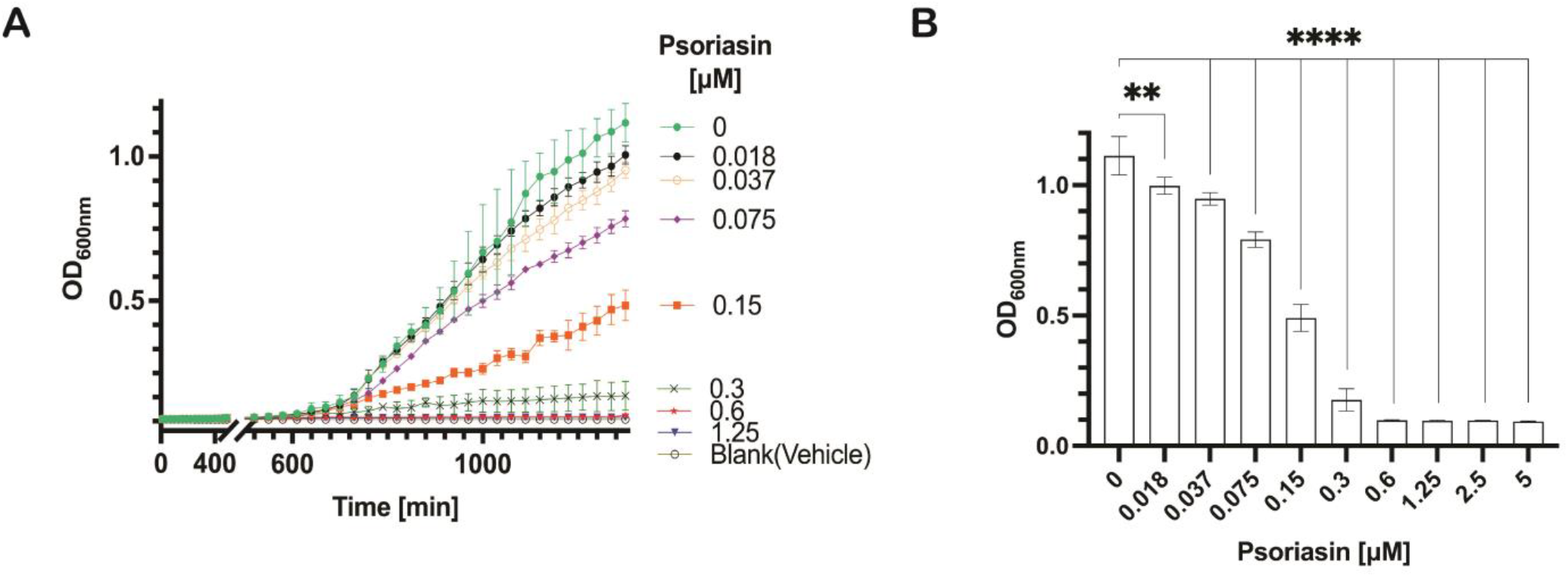
Growth curves of *C. albicans* in the presence of varying psoriasin concentrations. **(A)** Continuous monitoring of fungal growth in a 96-well plate was conducted by measuring absorbance at 600nm. Error bars represent the average of three replicates (±SD). **(B)** Bars showing the end-point absorbance values for each psoriasin concentration. Errors bars represent the average absorbance from three replicate wells (±SD). ^**^p < 0.005, ^****^p < 0.0001.

### 3.3 Cytotoxicity assessment of psoriasin on mammalian cells

Despite being a natural antifungal protein within the human saliva, we evaluated the cytotoxic effects of psoriasin on HeLa and human gingival fibroblast (HGF) cells. These two cell lines are classified as outer-layered epithelial and fibroblasts cells, respectively, and play a vital role in resisting pathogen invasion. The cells were treated with various psoriasin concentrations for 24 hours and their viability rates were evaluated using the Alamar blue assay, which measures metabolic activity as an indicator of cellular oxidation level and cytotoxicity. The results indicate that psoriasin does not impair cellular viability (Fig. 3), supporting its safe use in therapeutic applications.

**Figure 3.**
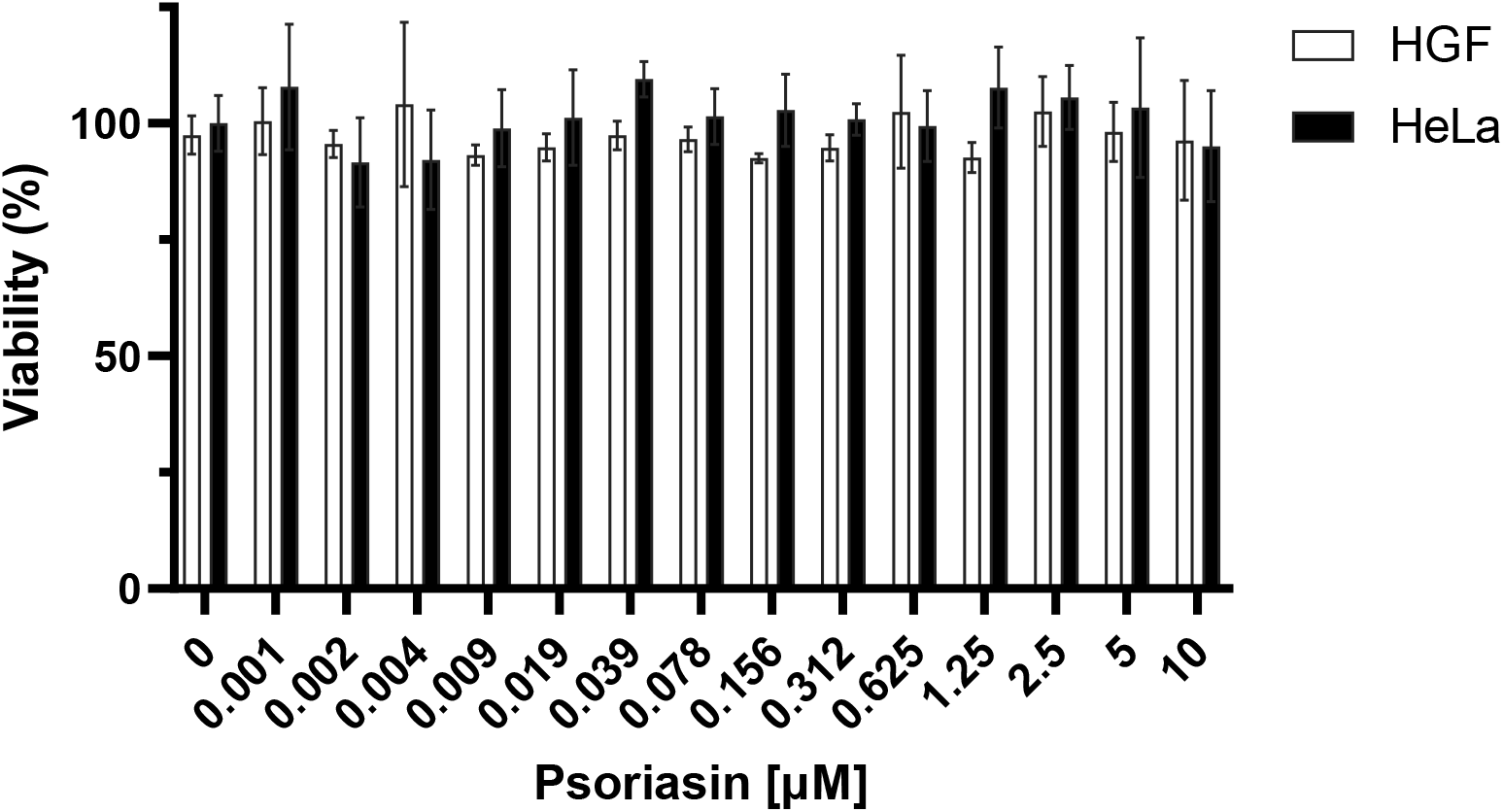
Effect of psoriasin on mammalian cellular viability. Alamar blue assay results show the viability of HeLa and HGF cells after 24 h incubation with variable concentrations of psoriasin. Results are shown as percentages relative to non-treated cells (left column). Errors bars represent the average signal originating from four replicate wells (±SD).

### 3.4 Psoriasin treatment on porous PMMA discs Inhibits *C. albicans* adhesion

To evaluate whether psoriasin could be effectively used as a preventive treatment on porous PMMA denture bases, we compared its activity to the effective anti-fungal agent voriconazole. PMMA discs were immersed in PBS solutions of either psoriasin (50 µM) or voriconazole (28 µM) for 24 hours. After thorough washing with PBS, the discs were exposed to a *C. albicans* culture in RPMI-MOPS medium and incubated at 30°C for 24 hours.

We then assessed the growth of *C. albicans* in the medium surrounding each disc (Fig. 4A) and evaluated the concentration of cells in the culture medium (Fig. 4B). The non-treated discs that were kept in RPMI without *C. albicans* (referred to as blank, top-left) showed no fungal growth, in contrast to the discs exposed to *C. albicans* (labeled as PBS, top-right). We observed minor fungal growth on the treated discs soaked with voriconazole (bottom-left) and psoriasin (bottom-right). Furthermore, we assessed the optical density (OD) at 600 nm for each well (Fig. 4B), which correlated with the data showing the absence of *C. albicans* growth in the medium containing the treated discs.

**Figure 4.**
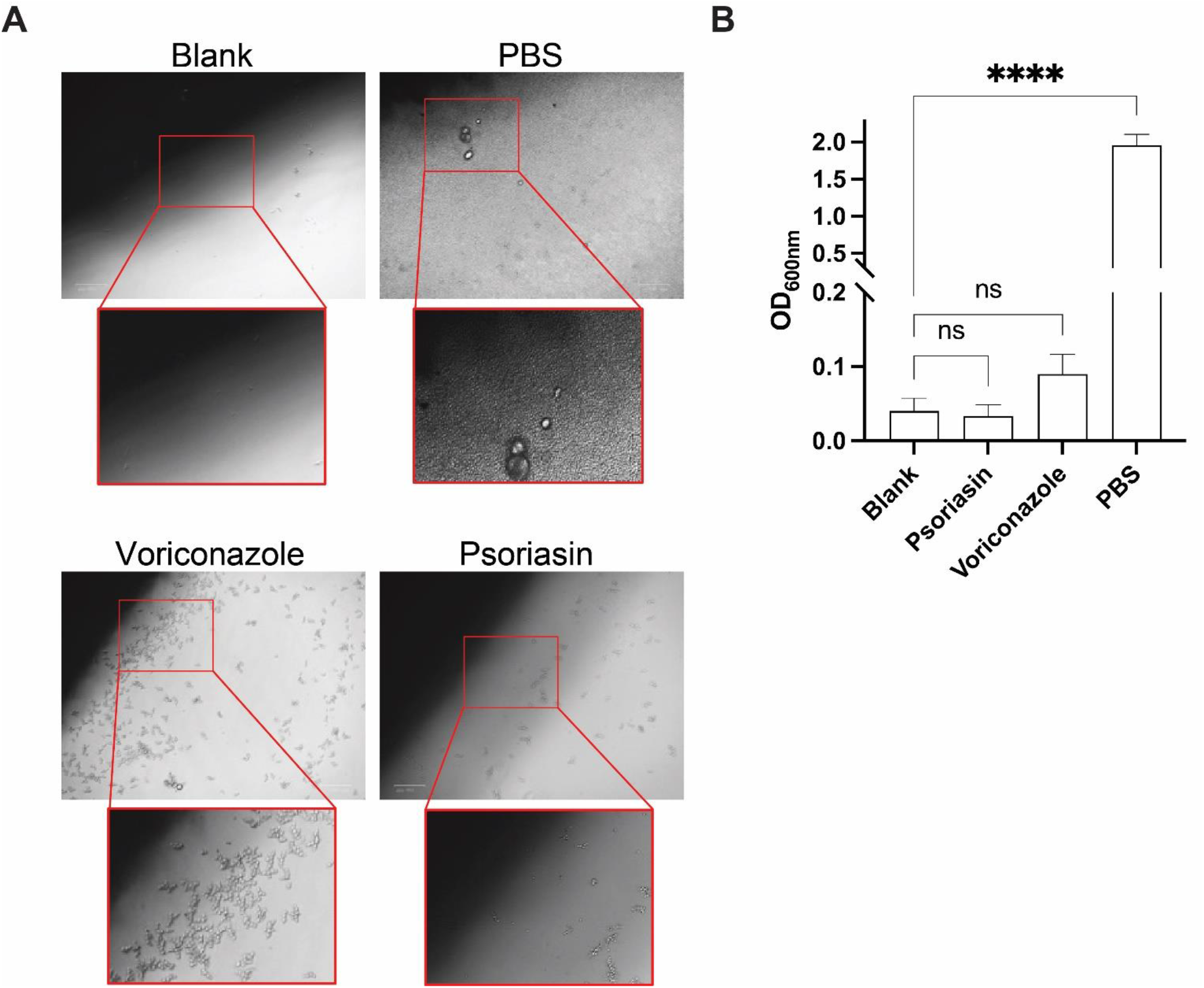
Effects of PMMA disc treatments on *C. albicans* growth. **(A)** PMMA discs were soaked with either psoriasin or voriconazole (50µM and 28 µM respectively) placed in a 24-well plate inoculated with *C. albicans* for 24 hours. Images display each disc following extensive washing with PBS. **(B)** Bar plot of the absorbance at 600 nm of the culture in each well following the 24-hour incubation. Errors bars represent the average absorbance of three replicate wells (±SD). ns -non significant; ^****^p < 0.0001.

### 3.5 SEM analysis reveals reduced *C. albicans* adhesion on psoriasin-treated PMMA discs

In addition to assessing the medium, we applied SEM to visually examine fungal adhesion and growth on the untreated and treated PMMA discs. The SEM images provided a clear contrast in *C. albicans* density between psoriasin-treated discs and those treated with PBS (Fig. 5A). Figure 5B shows the quantification of the fungal coverage area relative to the disc total area. Remarkably, the disc soaked in psoriasin solution shows better inhibition compared to voriconazole treated disc with 60% reduction in fungal growth of psoriasin relative to voriconazole treated discs. This effect was also evident in the RPMI medium results from discs pretreated with psoriasin (Fig. 4). This confirms the potential of psoriasin as a robust antifungal treatment for PMMA denture bases.

**Figure 5.**
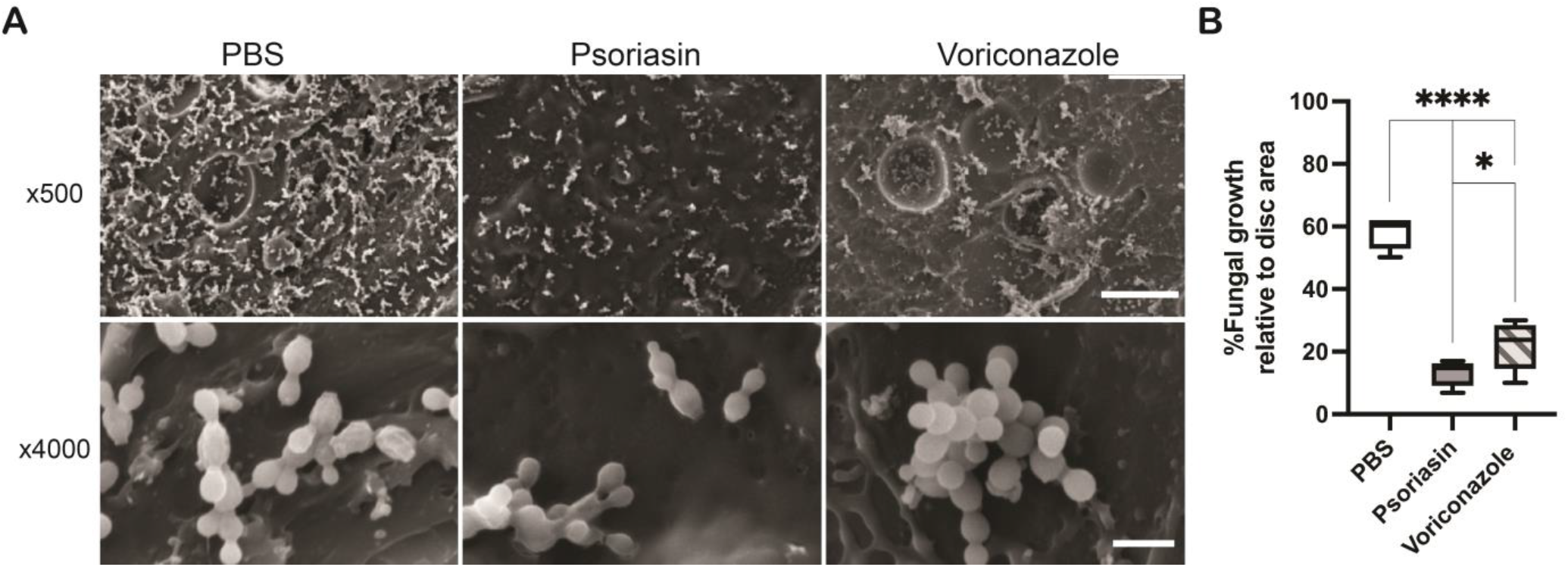
SEM analysis of *C. albicans* adhesion on PMMA discs soaked with psoriasin. **(A)** Representative SEM images with a magnification of x500 (top row) and x4000 (bottom row) following soaking of PMMA discs with PBS (left), 50 μM psoriasin (middle) and 28 μM voriconazole (right) and subsequently culturing of the discs in RPMI with *C. albicans*. Scale bar of top and bottom rows = 50 µm and 5µm, respectively. Measurement parameters 15 keV and 12 mm. **(B)** Quantification of percentage of fungal growth relative to disc area. Horizontal lines indicate the mean growth area of *C. albicans*. Values were measured using the Threshold tab in Fiji software and indicate the mean ± SD of five replicates. ^*^p < 0.05, ^****^p < 0.001.

## 4. Discussion

The prevalence of denture stomatitis has led to extensive search for innovative and effective strategies aimed at preventing microbial growth on PMMA denture bases. These include incorporating antimicrobial agents like silver or titanium oxide nanoparticles, which have shown effectiveness against Candida albicans(35-37). Alternative methods involve coating the PMMA surface with zwitterion or hydrophilic molecular chains (38,39). In addition, incorporating 5% zinc dimethacrylate(40), or much lower concentration of 0.045% copper nanoparticles(41) has proven effective in inhibiting *C. albicans* growth. Similarly, iron oxide nanoparticles combined with plant extracts have demonstrated antifungal properties(42). Alternative strategies like pulsed Nd:YAG laser irradiation show promise in biofilm inhibition, aligning with the efficacy of photodynamic therapy in fungal growth control. Alternative strategies like pulsed Nd:YAG laser irradiation show promise in biofilm inhibition, aligning with the efficacy of photodynamic therapy in fungal growth control(43-45). Other chemical agents are commonly used as denture cleaners (46-49). Despite these innovations, the complete prevention of Candida growth and biofilm removal from denture surfaces remains a complex challenge in dental medicine(30).

While biologic modalities are becoming part and offering a promising therapeutic avenue in various medical conditions, their application in dental treatments, including denture stomatitis, remains relatively unexplored with only a limited number of reported studies(32,50,51). Given the growing interest in natural proteins with antimicrobial properties in both medical and biotechnological fields and owing to psoriasin’s proven antibacterial and antifungal effectiveness(28,29), our study focused on evaluating psoriasin as a potential treatment option for denture stomatitis.

Through recombinant proteins expression and purification methods, we successfully produced high-yield and purity of psoriasin (Fig. 1), observing its strong inhibitory effect on *C. albicans* with an IC50 of 115 nM (Fig. 2). This is in contrast to psoriasin extracted from lesional psoriatic scale that did not show inhibitory effect against *C. albicans(52)*. Moreover, the recombinant psoriasin exhibited no adverse impact on mammalian cell viability, affirming its potential safety profile (Fig. 3). We then assessed its potential in preventing C. albicans colonization on porous PMMA. Pre-soaking PMMA discs in psoriasin solutions and then exposing them to *C. albicans*, we observed effective inhibition of fungal growth both in the culture medium and directly on the discs (Fig. 4-5). This demonstrates the effectiveness of psoriasin in preventing fungal colonization on PMMA surfaces.

We also note a few limitations of the current *in-vitro* study. While psoriasin naturally occurs in the oral cavity and is involved in various cellular processes, the impact of high concentrations in vivo could differ from in-vitro results and requires careful examination using suitable models. Additionally, the stability of psoriasin in the oral environment, with its diverse microbial activity, might affect its effectiveness and degradation rate. These challenges could potentially be addressed through protein engineering techniques, which could modify the protein structure to enhance stability and other properties(53).

Finally, it is enlightening to discuss psoriasin’s potential clinical applications and limitations in the context of denture stomatitis. This study’s findings indicate psoriasin efficacy as a treatment. Psoriasin could be used topically or as a disinfectant in clinical settings. It might either replace or complement current antifungal treatments, potentially allowing for reduced dosages of conventional medications. In addition, psoriasin solution could be used to store dentures when they are not in use, such as during sleep time. Given psoriasin positive effect on wound healing in keratinocyte cells(27), it could also help alleviate gum irritation and ulcerations often experienced by individuals suffer from denture stomatitis(54). Further research is needed to explore additional applications of psoriasin for various localized *Candida* infections and to other pathogens, including dermal and vaginal sites. Topical administration of antifungal proteins like psoriasin could present a viable treatment approach. These possibilities highlight the need for further research to fully understand and harness psoriasin’s potential in dental medicine and beyond.

## 5. Conclusions

There is an urgent need to develop additional antimicrobial solutions in general and dental medicine. Among these, denture stomatitis is a prevalent oral condition that is closely associated with *Candida* overgrowth within the oral mucosa, particularly beneath dentures. We describe here the application of psoriasin from a recombinant source to treat denture base for the prevention of fungal growth. The protein shows effective inhibition of *C. albicans* in medium and on the polymerized PMMA, marking it as a potential natural antimicrobial solution.

## Author Contributions

Lucia Adriana Lifshits, Edward Brohnshtein, May Attias, Yoav Breuer, Adi Cohen, Matan Gabay and Marina Sova, contributed to contributed to data acquisition, analysis, and interpretation and drafted the manuscript. Evgeny Weinberg, Eran Zenziper, Daniel Z. Bar, Nir Sterer and Maayan Gal contributed to the conception and design, contributed to data analysis and interpretation and drafted the manuscript. Daniel Z. Bar, Nir Sterer and Maayan Gal critically revised the manuscript. All authors gave final approval and agreed to be accountable for all aspects of the work ensuring integrity and accuracy.

## Acknowledgments

The research was supported by funding from Bountica Ltd and the School of Dental Medicine, Tel Aviv University and the Marian Gertner Institute for Medical Nano-Systems. L.A.L is grateful for Ph.D. scholarship financial support from the ADAMA Center for Novel Delivery Systems in Crop Protection. A.C is grateful for Ph.D. scholarship financial support from the Longevity and Health Research Center in Tel Aviv University.

