## Supplementary material for "Anti-fungal recombinant psoriasin effectively inhibits Candida albicans growth on denture base": Fig. S1

***Equal first-author contribution**

**Corresponding authors**

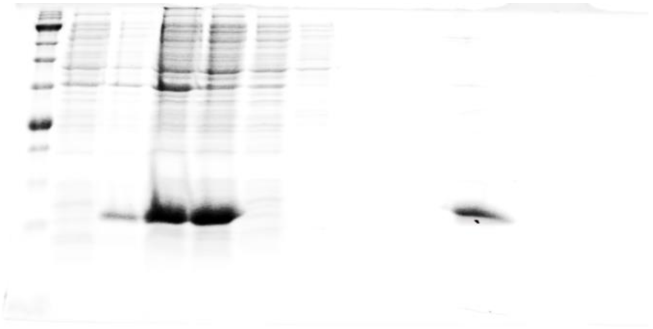

**Fig. S1. SDS-PAGE gel illustrating psoriasin purification.** Non-edited image of Fig. 1B.

**Supplementary Text S1**

**DNA sequence that was synthesized (Genscript Ltd.)**

ATGGGCAGCAGCCATCATCATCATCATCACAGCAGCGGCCTGGTGCCGCGCGGCAGCCA
TATGAGCAACACCCAGGCGGAGCGTAGCATCATTGGTATGATCGATATGTTCCACAAGTAC
ACCCGTCGTGACGATAAGATTGAGAAACCGAGCCTGCTGACCATGATGAAAGAAAACTTCC
CGAACTTTCTGAGCGCGTGCGATAAGAAAGGCACCAACTACCTGGCGGACGTGTTTGAGA
AGAAAGACAAGAACGAAGATAAGAAAATCGACTTCAGCGAATTTCTGAGCCTGCTGGGTGA
TATTGCGACCGACTATCACAAACAGAGCCACGGCGCGGCGCCGTGCAGCGGTGGCAGCC
AATAA

**Supplementary Text S2**

**Psoriasin protein sequence**

MGSSHHHHHHSSGLVPRGSHMSNTQAERSIIGMIDMFHKYTRRDDKIEKPSLLTMMKENFPNF
LSACDKKGTNYLADVFEKKDKNEDKKIDFSEFLSLLGDIATDYHKQSHGAAPCSGGSQ

HHHHHH - **His tag**
LVPRGS - **Thrombin cleavage site**
